# Testis morphogenesis and fetal Leydig cell development are mediated through ERK signaling activated by platelet derived growth factor receptor alpha

**DOI:** 10.1101/2023.07.21.549565

**Authors:** Shu-Yun Li, Satoko Matsuyama, Sarah Whiteside, Xiaowei Gu, Jonah Cool, Blanche Capel, Tony DeFalco

**Affiliations:** Reproductive Sciences Center, Division of Developmental Biology, Cincinnati Children’s Hospital Medical Center, Cincinnati, OH 45229 U.S.A.; Department of Cell Biology, Duke University Medical Center, Durham NC 27710 U.S.A.; Department of Pediatrics, University of Cincinnati College of Medicine, Cincinnati, OH 45267 U.S.A.

**Keywords:** *Pdgfra*, ERK, Leydig cells, testis cord, steroidogenesis.

## Abstract

*Platelet derived growth factor receptor alpha* (*Pdgfra*) plays a crucial role in the morphogenesis and differentiation of the fetal testis, but the molecular signaling involved in these processes remains unclear. Here, we use XY *Pdgfra*-null knock-in (KI) gonads to investigate the molecular mechanisms underlying *Pdgfra*-mediated testis cord formation and development of fetal Leydig cells (FLCs). The extracellular signal-regulated kinase (ERK) pathway, a well-known mitogen-activated protein kinase signaling pathway, was significantly inhibited in XY *Pdgfra* KI gonads, suggesting that ERK signaling is activated downstream of PDGFRA. Using *ex vivo* whole-organ culture, small interfering RNA (siRNA) cell culture methods, transwell assays and a genetic mouse model to disrupt ERK signaling in gonadal cells, we found that the ERK pathway promoted testis cord formation via *early growth response 1* (*Egr1*)-mediated cell migration and regulated the expression of steroidogenic enzymes in FLCs via activating the transcription factor cAMP responsive element binding protein 1 (CREB; official name CREB1). These findings highlight the significance of PDGFRA and its downstream signaling in fetal testis morphogenesis and cellular differentiation, thus providing insights into the etiology of and potential therapeutic targets for disorders related to gonadal development.

## Introduction

Platelet derived growth factor receptor alpha (PDGFRA) belongs to a family of receptor tyrosine kinases (RTKs), whose activation via binding of PDGF ligands regulates multiple downstream pathways within the cell, including the extracellular signal-regulated kinase (ERK) and mitogen-activated protein kinase (MAPK) pathways^1^. The PDGF ligands and PDGF receptors (PDGFRs), PDGFRA and PDGFRB, play critical roles during development by driving cellular responses including proliferation, survival, migration, and differentiation^2^. Knockout mouse studies have indicated that PDGFB and PDGFRB are required for the development of support cells in the vasculature, whereas PDGFA and PDGFRA are more broadly essential during embryogenesis^3–8^. In the fetal testis, *Pdgfra* is mainly expressed in interstitial mesenchymal cells and its ligands are also expressed in the fetal testis: *Pdgfa* is strongly expressed in Sertoli cells within testis cords; *Pdgfb* is expressed in endothelial cells; and *Pdgfc* is expressed at the gonad/mesonephros boundary^9^. PDGFRA acts as a key player in the developmental program downstream of *Sry* that orchestrates the morphological and cellular development of the early testis, including vascular patterning, testis cord formation, sex-specific cell proliferation, endothelial cell migration, and fetal Leydig cell (FLC) differentiation^9^. PDGF signaling can trigger several molecular cascades, such as the MAPK and phosphoinositide-3-kinase/thymoma viral proto-oncogene 1 (PI3K/AKT) pathways, and can induce the activation of several critical factors, such as early growth response 1 (EGR1). EGR1, a mammalian transcription factor, is important for cell migration of multiple cell types, including endothelial cells and mammary gland cells^10,11^, and it is induced by PDGF in human pulmonary artery smooth muscle cells (hPASMCs) *in vitro*^12^. However, the underlying molecular mechanisms of how *Pdgfra* promotes morphological and cellular development of the fetal testis are poorly defined.

The MAPK pathway regulates a variety of biological processes such as cell proliferation, differentiation, and apoptosis. It is also involved in numerous male reproductive processes, including spermatogenesis, sperm maturation and activation, capacitation, and the acrosome reaction^13^. ERK, a type of serine/threonine protein kinase whose members include ERK1 (official name MAPK3) and ERK2 (official name MAPK1), belongs to the MAPK family and is required for proliferation, differentiation, survival, migration, and apoptosis during vertebrate embryonic development^14–16^. In the adult testis, testosterone-mediated ERK signaling facilitates the adhesion of immature germ cells to Sertoli cells^17^, and depletion of MEK/ERK signaling results in a reduction of functional adult Leydig cells (ALCs)^18^. Despite a number of studies demonstrating the importance of the ERK pathway in male reproduction, the expression pattern of activated ERK and its regulation in the fetal testis are not well understood.

Seminiferous tubules, the site of spermatogenesis where germ cells develop into spermatozoa in close interaction with Sertoli cells, are the fundamental units of testis structure and function. Testis cords are embryonic precursors of the seminiferous tubules, and disruption of testis cord formation results in gonadal dysgenesis, thus impairing male fertility^19, 20^. Testis cord formation is hindered in *Pdgfra*-null XY gonads, due at least in part to a failure of mesonephric cell migration into the gonad^9^. However, the signaling pathways involved in *Pdgfra*-mediated testis cord formation are not well characterized.

Leydig cells (LCs) are steroidogenic cells present in the testicular interstitial compartment adjacent to the seminiferous tubules. They are the primary source of androgens in males, which are required for sexual differentiation and spermatogenesis throughout fetal and adult life^21^. In rodents, there are two distinct Leydig cell populations, fetal (FLCs) and adult (ALCs). In mice, the FLC population begins to arise shortly after sex determination at E12.5, peaks shortly before birth, and declines over the first 2 weeks of postnatal life^22^. FLCs only produce precursor androgens, such as androstenedione, which is converted to testosterone by 17β-hydroxysteroid dehydrogenase type 3 (HSD17B3) expressed in fetal Sertoli cells; since ALCs express HSD17B3 postnatally, these cells can produce testosterone on their own^23, 24^. FLCs arise via differentiation of FLC progenitors or stem cells in the testicular interstitium, which originate from distinct cellular sources such as the coelomic epithelium (CE) and perivascular cells at the gonad–mesonephros border^25, 26^. A single-cell RNA-seq study revealed that *Sf1*-positive cells (official name *Nr5a1*) are interstitial progenitors of FLCs^27^, and Leydig cell defects in *Pdgfra*-knockout testes might be due to reduced coelomic epithelial proliferation^9^. *Sphingosine phosphate lyase 1* (*Spgl1*) and *pleckstrin homology domain– containing family A1* (*Plekha1*) are 2 putative downstream targets of PDGF signaling that are required for the LC population in postnatal and adult testes^28^. However, testicular morphology and male development appear normal before postnatal day 20 in both *Sgpl1*- and *Plekha1* deficient mice, indicating that these 2 genes are unlikely to be involved in fetal testis morphogenesis or differentiation^28^.

The mechanisms of how *Pdgfra* regulates cell migration, testis cord morphogenesis, and FLC development remain unclear. Here we investigate the molecular pathways underlying testicular defects observed in *Pdgfra*-deficient embryos. Using a *Pdgfra*-GFP knock-in (KI) allele, we found that deletion of *Pdgfra* resulted in inactivation of ERK signaling throughout the fetal testis. Using *ex vivo* and *in vitro* experiments, we show that ERK regulates testis cord formation via the transcriptional regulator early growth response 1 (EGR1). While we observed a mild disruption in sex-specific vasculature, which is required for the maintenance of perivascular FLC progenitors^26^, in *Pdgfra*-null gonads, we did not see any sustained reduction of FLC progenitors. However, we did detect a reduction in the mRNA and protein levels of terminal LC markers (e.g., *Cyp11a1*, *Hsd3b1*, and *Cyp17a1*), despite a normal number of typical NR5A1-expressing cells in XY *Pdgfra*-knockout gonads, suggesting that the progenitors could initially give rise to FLCs but lost their downstream steroidogenic function. We conditionally deleted ERK signaling in *Nr5a1*-expressing cells in XY gonads using *Nr5a1*-Cre; *Mapk1*^flox/flox^; *Mapk3^-/-^* mice and found no effect on progenitors or initial FLC specification, but observed a significant and specific disruption of the steroidogenic program. Our work provides new insights into the mechanisms underlying of the role of *Pdgfra* in testis cord formation and FLC development via the ERK pathway in the fetal testis.

## Results

### *Pdgfra* is required for testis cord formation and Leydig cell development

Given that previous analysis of *Pdgfra* utilized a traditional knockout allele, we first assessed the testis phenotype of *Pdgfra* null mutation caused by a GFP knock-in (KI) allele^29^ (Fig. S1). Our analyses revealed that *Pdgfra*^GFP/GFP^ XY KI gonads recapitulated testicular defects similar to the previously described null phenotype^9^. The formation of testis cords was severely disrupted at E12.5 (Fig. 1A) and at E13.5 resulted in large, looping cords in 40% fewer numbers in *Pdgfra*^GFP/GFP^ XY KI gonads as compared to controls (Fig. 1C, 1E). qRT-PCR results revealed that mRNA levels of the Sertoli cell marker *Sox9* were not affected in homozygous XY KI gonads (Fig. 1F), suggesting that Sertoli cell identity and number were normal. As expected, based on previous studies, CYP11A1 (a FLC marker) expression was significantly reduced in E12.5 (Fig. 1B) and E13.5 (Fig. 1D) *Pdgfra*^GFP/GFP^ XY KI gonads. Consistent with immunofluorescence data for CYP11A1, mRNA levels of the FLC markers *Cyp11a1*, *Hsd3b1*, and *Cyp17a1* were all significantly reduced in E12.5 and E13.5 *Pdgfra*^GFP/GFP^ XY KI gonads (Fig. 1F). Consistent with a previous report using a traditional knockout allele^9^, *Pdgfra*^GFP/GFP^ XY KI gonads possessed a grossly normal coelomic arterial vessel at E12.5 (Fig. 1A) and E13.5 (Fig. 1C), even though vasculature failed to promote normal testis cord formation. Using qRT-PCR, we found that mRNA expression of the endothelial marker *Cdh5* was significantly decreased in E12.5 and E13.5 *Pdgfra*^GFP/GFP^ XY KI gonads (Fig. 1F), although not to the extent of steroidogenic genes. Later in fetal development and consistent with earlier stages, E15.5 XY KI gonads showed aberrant testis cord architecture and a severe decrease in Leydig cell number (Fig. 1G).

**Figure 1.**
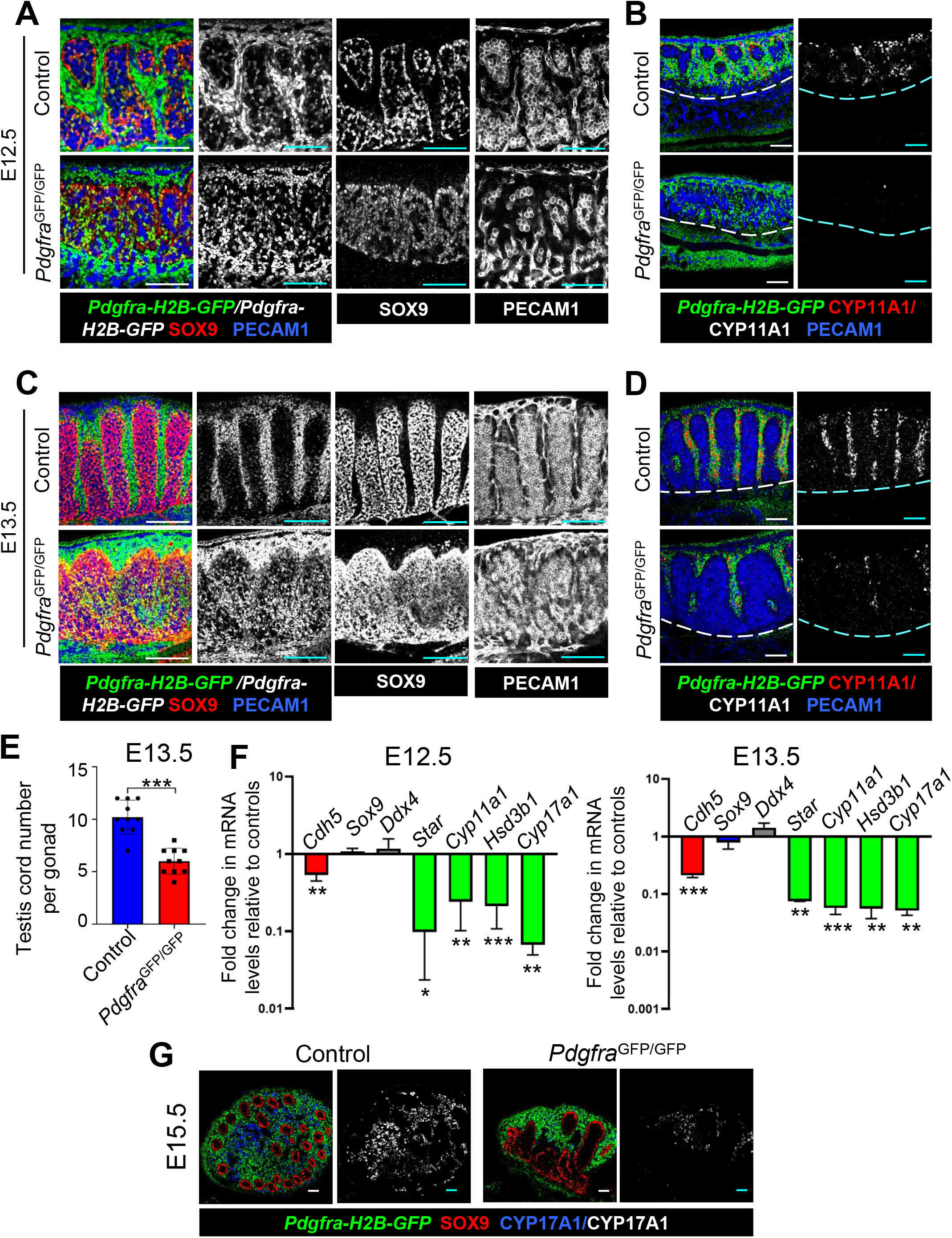
*Pdgfra* is required for testis development. (A-D) Immunofluorescence images of E12.5 (A,B) and E13.5 (C,D) control (*Pdgfra*^GFP/+^) and *Pdgfra*^GFP/GFP^ XY KI gonads. (E) Quantification of testis cord number in E13.5 control and *Pdgfra*^GFP/GFP^ XY KI gonads. (F) qRT-PCR analyses of E12.5 and E13.5 control and *Pdgfra*^GFP/GFP^ XY KI gonads. (G) Immunofluorescence images of control and E15.5 *Pdgfra*^GFP/GFP^ XY KI gonads. Scale bar: 100 μm. \**P*<0.05; \*\**P*<0.01; \*\*\**P*<0.001.

Sexually dimorphic vascular remodeling is one of the earliest steps during sex-specific morphogenesis and directs testis cord formation^30^. Previous recombinant gonad culture experiments using *Pdgfra* knockout gonads revealed that *Pdgfra* is required to induce the migration of mesonephric cells, such as endothelial cells, into the gonad^9^. Our live imaging of *Kdr*-myr-mCherry; *Pdgfra*-GFP XY gonads, which are simultaneously labeled for endothelial and mesenchymal cells, initiated at E11.75 showed that control XY gonads quickly form a coelomic vessel and induce the aggregation of GFP-positive mesenchyme around microvasculature. In contrast, *Pdgfra*^GFP/GFP^ XY KI gonads still recruited endothelial cells and formed a coelomic vessel, but formation of the vessel was not apparent until the end of the imaging time period (Fig. S2A), suggesting a delay of vascularization in *Pdgfra*^GFP/GFP^ XY KI gonads. PECAM1 immunofluorescence showed that in *Pdgfra*^GFP/GFP^ XY KI gonads the coelomic vessel formed fewer distinct branches between cords as compared to controls (Fig. S2B), indicating that a delay in vascularization may impair the formation of testis cords. Taken together, our data suggest that the reduction in mesonephric cell migration results in the delay of vascularization in *Pdgfra-*null XY gonads.

### Inactivation of the ERK pathway impairs testis cord formation in *Pdgfra*^GFP/GFP^ XY KI gonads

The ERK pathway is a well-known signaling pathway downstream of PDGF that is involved in multiple cellular and developmental responses^31^, but the role(s) and expression pattern of p-ERK (phosphorylated-ERK, which represents its activated form) in early fetal testes are unclear. Here, we observed that p-ERK was widely expressed in Sertoli cells, interstitial cells, and endothelial cells in E13.5 XY controls, while it was significantly decreased throughout *Pdgfra*^GFP/GFP^ XY KI gonads (Fig. 2A). To examine the roles of p-ERK in testis development, we treated E11.5 XY gonads *ex vivo* with U0126, which is an inhibitor of MEK/ERK kinases^32^. After 48 hours of *ex vivo* culture with U0126, p-ERK staining was dramatically reduced (Fig. 2B), thus demonstrating that the ERK pathway was inactivated. Compared to controls in which SOX9+ Sertoli cells were regularly spaced to form testis cords, SOX9+ Sertoli cells were uniformly distributed in XY gonads treated with U0126 (Fig. 2C). While U0126 treatment significantly inhibited the ERK pathway, it did not disrupt testis cord formation when gonads were cultured starting at E12.5 (Fig. S3A and B). PECAM1 immunofluorescence analyses also showed a reduction in coelomic vessel formation in U0126-treated samples (Fig. 2C, 2D and S3A-C). However, impaired vasculature did not significantly affect FLC number (Fig. 2D and S3C). qRT-PCR analysis revealed a dramatic reduction of *Cdh5* mRNA levels in E11.5 and E12.5 U0126-cultured XY gonads, while Sertoli cells (*Sox9*), germ cells (*Ddx4*), and FLCs (*Star*, *Cyp11a1*, *Hsd3b1*, and *Cyp17a1*) did not significantly change (Fig. 2E and S3D). These results suggest that the inhibition of ERK signaling impairs testis cord formation via the disruption of vascularization at E11.5. However, once cord structure is already established at E12.5, the inactivation of ERK is unable to disturb this process.

**Figure 2.**
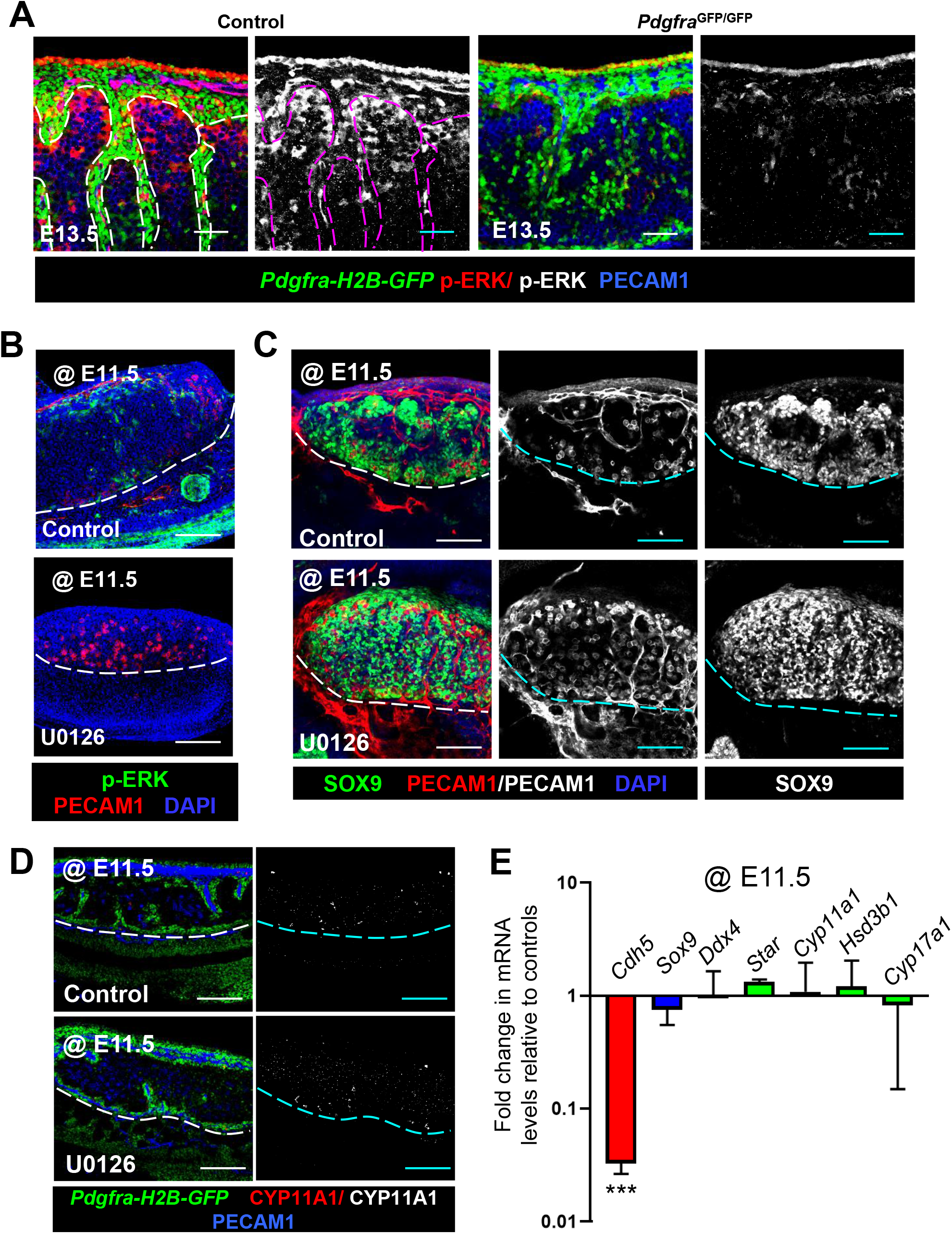
Inactivation of p-ERK impairs testis cord formation in E11.5 XY gonads. (A) Immunofluorescence images showing p-ERK expression pattern in control (*Pdgfra*^GFP/+^) and E13.5 *Pdgfra*^GFP/GFP^ XY KI gonads. (B-E) Immunofluorescence images (B-D) and qRT-PCR analyses (E) of E11.5 XY gonads cultured with U0126 for 48 hours *ex vivo*. Scale bar: 100 μm. \*\*\**P*<0.001.

### EGR1 mediates cell migration and is downregulated in *Pdgfra*^GFP/GFP^ XY KI gonads

We found that both mRNA and protein levels of *Egr1* were significantly reduced in *Pdgfra*^GFP/GFP^ XY KI gonads compared to controls (Fig. 3A and B). To determine whether *Egr1* regulates cell migration in fetal testes, a transwell migration assay was performed using media containing 10% fetal bovine serum (FBS) as a chemoattractant. Gonadal or mesonephric cells were isolated from E12.5 wild type XY gonad-mesonephric complexes, and each population was transfected with *Egr1* siRNA or a scrambled control siRNA. Forty eight hours after transfection, mRNA levels of *Egr1* were significantly down-regulated in both gonadal and mesonephric cells (Fig. 3C). Compared to control cells transfected with a scrambled siRNA, *Egr1* knockdown cells showed significantly reduced cell migration toward the chemoattractant (Fig. 3D and E). qRT-PCR analyses showed that endothelial cell (*Cdh5*), Sertoli cell (*Sox9*), and interstitial cell (*Nr2f2* and *Pdgfra*) gene expression did not significantly change (Fig. 3F), indicating that *Egr1* knockdown did not affect the identity of these cells. These data suggest that *Egr1* promotes gonadal and mesonephric cell migration in early fetal XY gonads.

**Figure 3.**
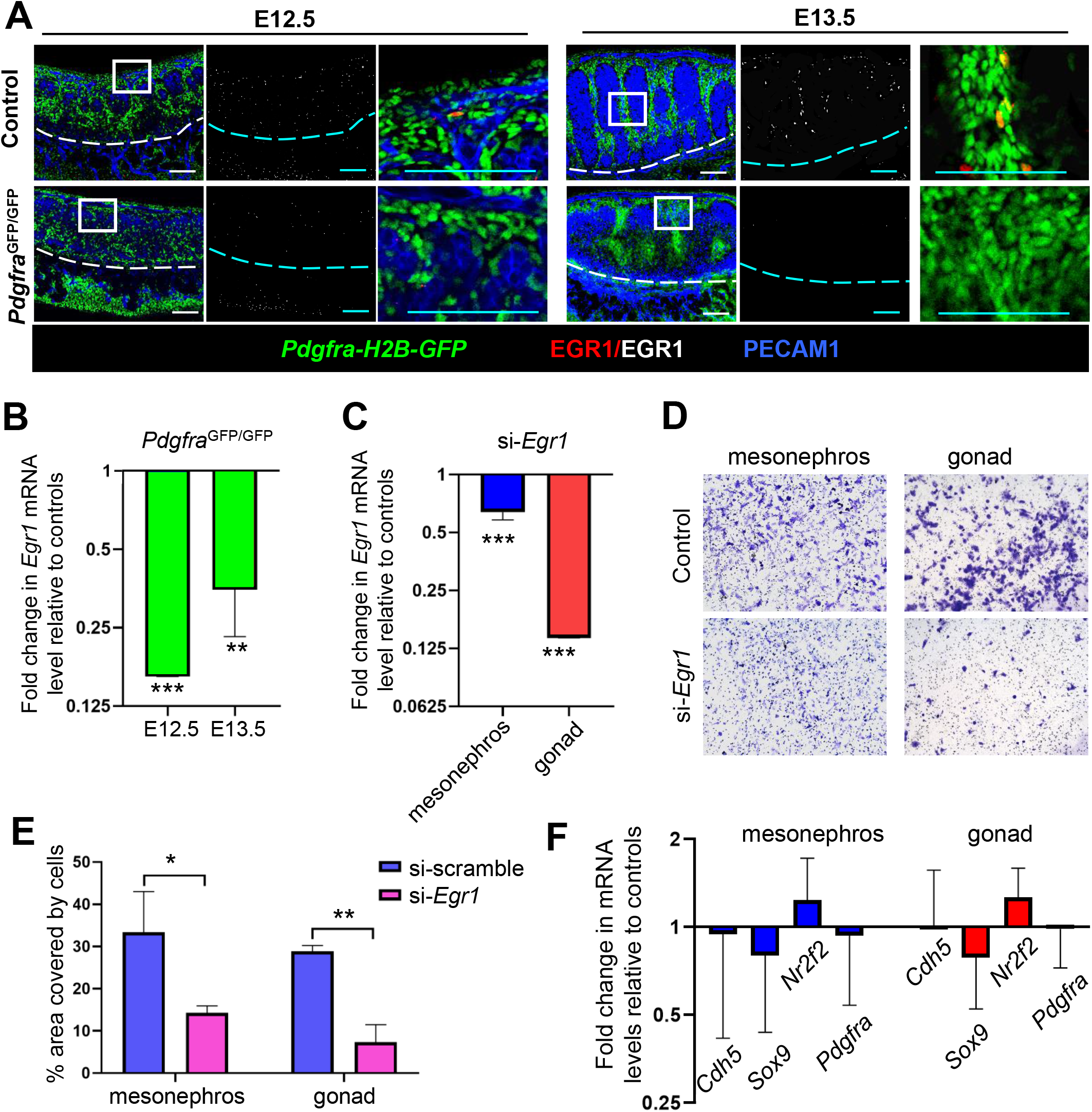
*Egr1* is involved in testis cord formation in E11.5 XY gonads. (A,B) Immunofluorescence images (A) and qRT-PCR analyses (B) showing EGR1 protein and *Egr1* mRNA levels in E12.5 and E13.5 control and *Pdgfra*^GFP/GFP^ XY KI gonads. (C-F) qRT-PCR analyses (C,F) and transwell cell migration assays (D,E) of gonadal and mesonephric cells transfected with control scrambled siRNA (si-scramble) or *Egr1* knockdown siRNA (si-*Egr1*). Scale bar: 100 μm. \**P*<0.05; \*\**P*<0.01; \*\*\**P*<0.001.

To examine whether ERK signaling regulates *Egr1*, we examined mRNA and protein levels of *Egr1* in E11.5 XY gonads cultured with U0126 for 48 hours, and we observed that they were significantly down-regulated compared to controls (Fig. 4A and B), indicating that ERK activation is required for expression of *Egr1* in XY gonads. We also found that U0126 significantly reduced mRNA expression of *Egr1* in gonadal and mesonephric cells *in vitro* (Fig. 4C). Furthermore, transwell assays revealed that U0126 treatment led to reduced cell migration of gonadal and mesonephric cells (Fig. 4D and E). qRT-PCR analyses showed that U0126 treatment had no effect on endothelial cell (*Cdh5*), Sertoli cell (*Sox9*), or interstitial cell (*Nr2f2* and *Pdgfra*) gene expression (Fig. 4F). Taken together, our data suggest that inactivation of ERK signaling impairs migration of gonadal and mesonephric cells via the suppression of EGR1.

**Figure 4.**
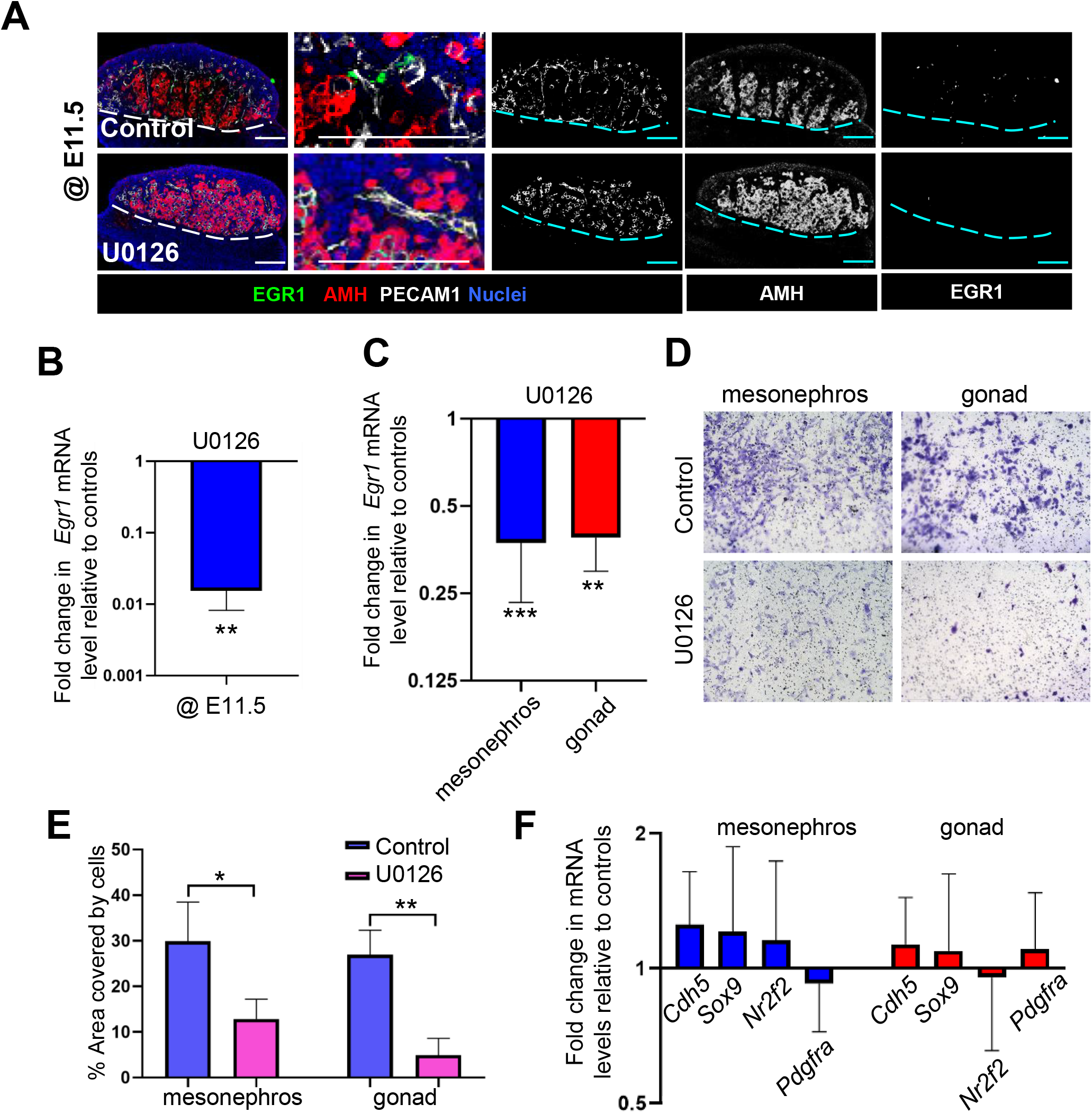
*Egr1* is regulated by ERK in E11.5 XY gonads. (A,B) Immunofluorescence images (A) and qRT-PCR analyses (B) of E11.5 XY gonads cultured with U0126 for 48 hours *ex vivo*. (C-F) qRT-PCR analyses (C,F) and transwell cell migration assays (D,E) of gonadal and mesonephric cells cultured with DMSO control or U0126 *in vitro*. Scale bar: 100 μm. \**P*<0.05; \*\**P*<0.01; \*\*\**P*<0.001.

### Knockout of *Pdgfra* specifically impairs the terminal steroidogenic program in FLCs

A previous study reported a disruption of sex-specific proliferation in *Pdgfra*-null XY gonads^9^, suggesting that this defect might be responsible for the reduction of FLCs. To examine the whether the reduction of FLCs in *Pdgfra*^GFP/GFP^ XY KI gonads was due to insufficient LC progenitors, we assessed the expression of the LC progenitor markers JAG1, Nestin, and NR2F2 using immunofluorescence. Levels of JAG1, Nestin, and NR2F2 were decreased in E12.5 *Pdgfra*^GFP/GFP^ XY KI gonads (Fig. 5A). qRT-PCR data also showed a significant reduction of mRNA levels of the FLC progenitor markers *Arx*, *Jag1*, *Nestin*, and *Nr2f2* in E12.5 KI gonads (Fig. 5B). We next examined the expression of these progenitor markers in E13.5 *Pdgfra* XY KI gonads. However, immunofluorescence images and qRT-PCR data indicated that protein and mRNA levels of these progenitor markers were back to levels similar to controls (Fig. 5A and B). To assess the proliferative status of the coelomic domain, which is comprised of the coelomic epithelium and subjacent mesenchyme, at E13.5, we examined the expression of the mitotic marker phosphorylated histone H3 (pHH3). Proliferation in both E13.5 control and *Pdgfra*^GFP/GFP^ XY KI gonads was similarly enriched in somatic cells throughout the coelomic domain (Fig. 5C). These results indicated that cells of the coelomic domain could proliferate at later stages; therefore, reduced proliferation was unlikely the underlying cause of FLC reduction in KI gonads.

**Figure 5.**
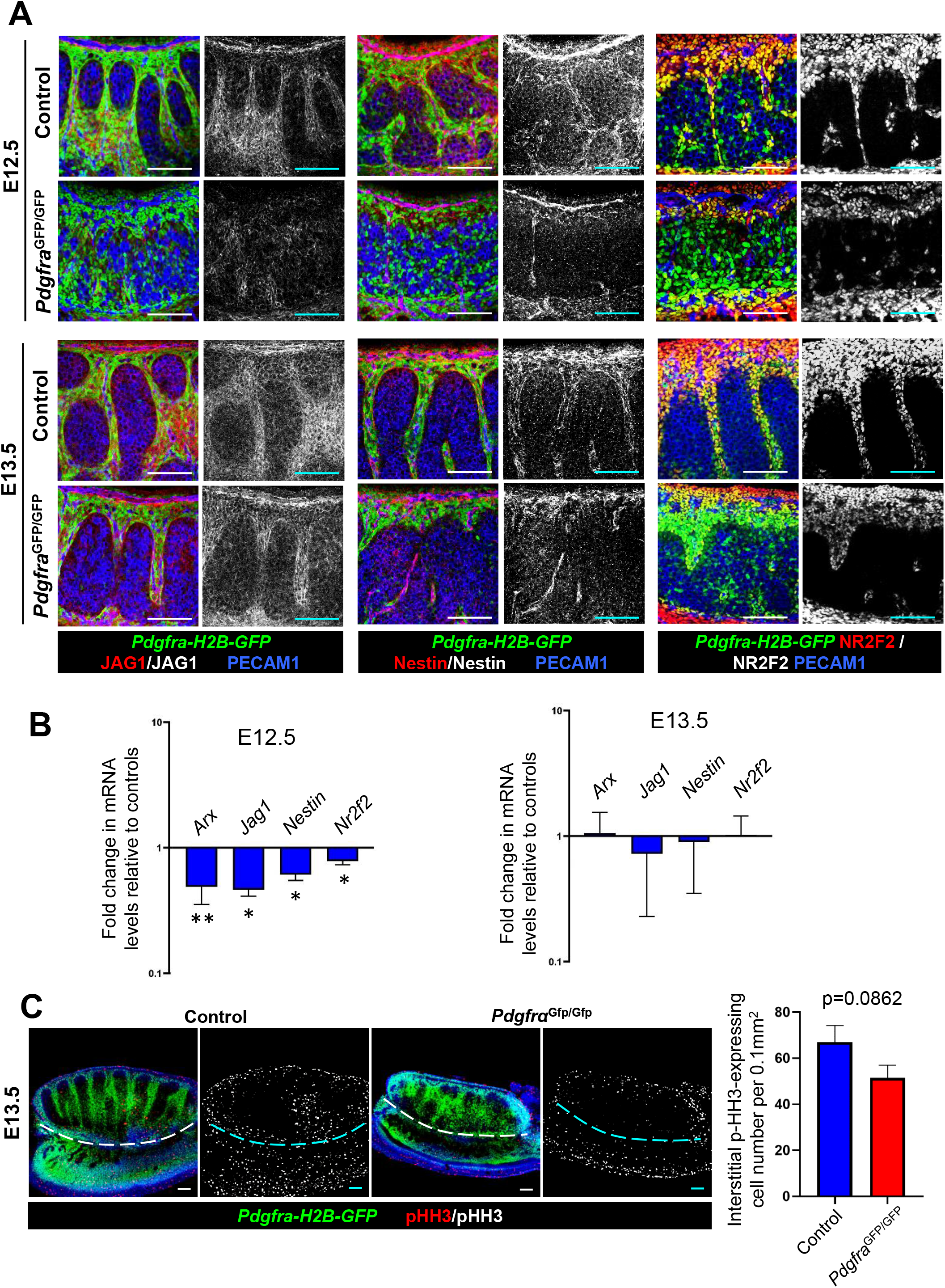
Interstitial LC progenitors are initially reduced, but quickly recover, in *Pdgfra*^GFP/GFP^ XY KI gonads. (A-C) Immunofluorescence images (A,C) and qRT-PCR analyses (B) of control (*Pdgfra*^GFP/+^) and *Pdgfra*^GFP/GFP^ XY KI gonads. Graph in C shows quantification of pHH3+ cells per unit area in E13.5 control and *Pdgfra*^GFP/GFP^ XY KI gonads. Scale bar: 100 μm. \**P*<0.05; \*\**P*<0.01; \*\*\**P*<0.001.

Despite the apparent recovery of FLC progenitors back to normal levels in KI gonads, it is still possible that they failed to differentiate into FLCs. To examine whether the differentiation of FLCs was disrupted, we analyzed the expression of the LC progenitor marker NR2F2 and the committed LC marker NR5A1 (also called SF1 or AD4BP) in E13.5 XY control and *Pdgfra* KI gonads. In E13.5 control and *Pdgfra*^GFP/GFP^ XY KI gonads, LC progenitors were identified via strong expression of NR2F2 and lack of NR5A1 expression, while LCs were identifiable by their large, round nuclei that were strongly positive for NR5A1 and negative for NR2F2 (Fig. 6A). In addition to these two clearly defined cell types, we also observed cells that weakly expressed both NR2F2 and NR5A1, suggesting these cells were undergoing the transition from NR2F2+ progenitors to NR5A1+ FLCs. Quantification of NR5A1+ FLCs showed that there was no significant difference between controls and *Pdgfra*^GFP/GFP^ XY KI gonads (Fig. 6B), suggesting that FLC progenitors successfully transitioned to the FLC differentiation program in E13.5 *Pdgfra*^GFP/GFP^ XY KI gonads. However, immunofluorescence images revealed that there was a significant increase in the number of interstitial NR5A1+ cells that did not co-express CYP11A1 in E13.5 and E15.5 *Pdgfra*^GFP/GFP^ XY KI gonads (Fig. 6C and F). Furthermore, cells in KI gonads that did express CYP11A1 expressed it more weakly than in controls, which is consistent with our whole gonad qRT-PCR data (Fig. 1). Cell number quantification confirmed that the number and percentage of NR5A1+ cells co-expressing CYP11A1 were significantly decreased in E13.5 *Pdgfra*^GFP/GFP^ XY KI gonads (Fig. 6D and E). Taken together, these data indicate that *Pdgfra* is not required for the initiation of FLC differentiation, but instead is specifically required for the downstream, terminal steroidogenic program in FLCs during fetal testis development.

**Figure 6.**
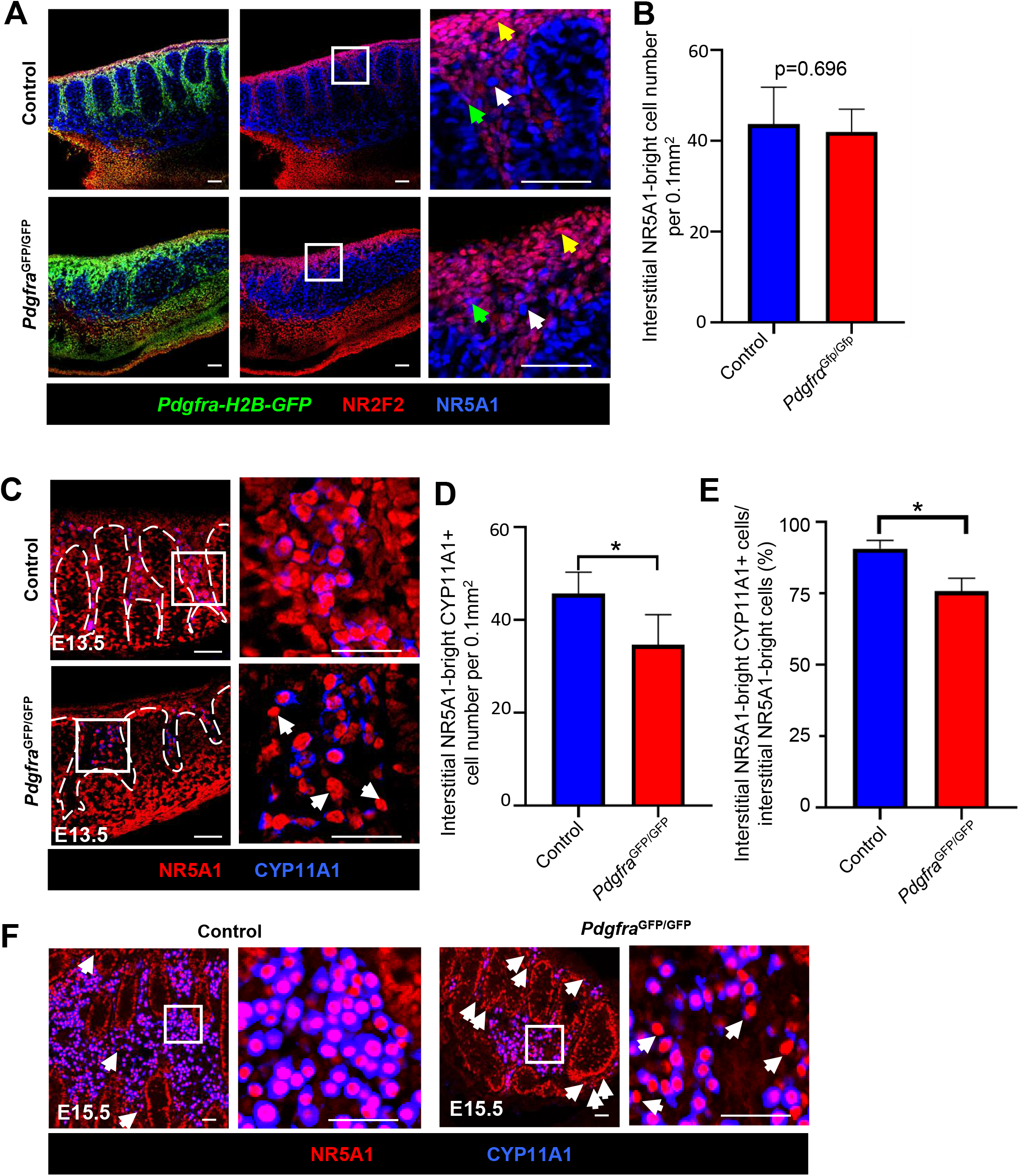
Initial specification of LC progenitors is normal in *Pdgfra*^GFP/GFP^ XY KI gonads. (A-C) Immunofluorescence images (A,C) of E13.5 control (*Pdgfra*^GFP/+^) and *Pdgfra*^GFP/GFP^ XY KI gonads and quantification of NR5A1+ FLCs (B) and NR5A1+ and CYP11A1+ double positive FLCs (D,E) in E13.5 control and *Pdgfra*^GFP/GFP^ XY KI gonads. (F) Immunofluorescence images of E15.5 control and *Pdgfra*^GFP/GFP^ XY KI gonads. Scale bar: 100 μm. \**P*<0.05.

### *Pdgfra* regulates expression of steroidogenesis enzymes in fetal Leydig cells via the ERK pathway

The MEK/ERK signaling cascade in LCs is important for the maintenance of functional LCs in the adult testis^18^, but its role in the fetal testis is unclear. Considering that p-ERK was significantly decreased in interstitial cells of *Pdgfra-*null XY gonads (Fig. 2A), we conditionally deleted the ERK pathway in gonadal interstitial cells using a global KO of *Erk1* (*Mapk3*^−/−^) and a conditional allele of *Erk2* (*Mapk1^flox/flox^*) deleted specifically in gonadal cells using *Nr5a1*-Cre mice. To verify the pattern of *Nr5a1*-Cre activity, we examined E13.5 *Nr5a1*-Cre; *Rosa*-Tomato XY gonads and we observed most Tomato+ interstitial and Sertoli cells expressed NR5A1 (Fig. S4A), demonstrating that we could target fetal gonadal cells effectively using *Nr5a1*-Cre.

In E14.5 control XY gonads (*Nr5a1*-Cre; *Mapk1^flox/+^*; *Mapk3*^+/-^), p-ERK was strongly expressed in vasculature, interstitium, and Sertoli cells, while it was robustly inactivated in interstitial and Sertoli cells in E14.5 double knockout (dKO) XY gonads (*Nr5a1*-Cre; *Mapk1^flox/flox^*; *Mapk3*^−/−^) (Fig. 7A). In dKO gonads, p-ERK only remained detectable in endothelial cells and faintly in germ cells (Fig. 7A), both of which are not targeted by *Nr5a1*- Cre. In E14.5 single knockout (sKO) XY gonads (*Nr5a1*-Cre; *Mapk1^flox/+^*; *Mapk3*^−/−^ or *Nr5a1*- Cre; *Mapk1^flox/flox^*; *Mapk3*^+/-^), intensity of p-ERK expression was also reduced compared to controls (Fig. S5A). Testis cords and vasculature in E14.5 dKO and sKO XY gonads were similar to controls (Fig. 7A and S5A). However, we observed a severe reduction of steroidogenic enzyme expression of StAR and CYP17A1 in E13.5 (Fig. S4B) and E14.5 dKO (Fig. 7B) gonads, while there was no detectable change in E14.5 sKO XY gonads (Fig. S5B). qRT-PCR analyses showed that endothelial cell (*Cdh5*), Sertoli cell (*Sox9*), germ cell (*Ddx4*), LC progenitor (*Lhx9, Arx, Nestin, Jag1,* and *Nr2f2*), and committed LC (*Insl3* and *Ren1*, which are non-steroidogenic markers of LCs) gene expression were not significantly decreased, while steroidogenesis enzymes (*Star, Cyp11a1, Hsd3b1,* and *Cyp17a1*) were all significantly reduced in E14.5 dKO XY gonads as compared to controls (Fig. 7C). We next sought to determine how the ERK pathway regulates steroidogenic enzyme expression in LCs. We performed genome-wide analyses for predicted transcription factor binding sites in the JASPAR CORE collection, focusing on the genomic regions near *StAR*, *Cyp11a1*, *Hsd3b1*, and *Cyp17a1*. We found that they all contained binding sites for cAMP responsive element binding protein 1 (CREB; officially known as CREB1), which is a nuclear transcription factor (Fig. S4C and Supplementary Table S1). In controls, p-CREB (activated CREB) was mainly expressed in GFP+ interstitial cells, while it was weakly expressed and in fewer cells in E13.5 *Pdgfra*^GFP/GFP^ XY KI gonads (Fig. 7D), suggesting that disruption of PDGF signaling restricts the activation of CREB. We next examined the expression pattern of p-CREB in ERK-depleted XY gonads and found that it was significantly reduced in E13.5 dKO XY gonads relative to controls (Fig. 7E). Taken together, our data suggests that *Pdgfra* regulates expression of steroidogenic enzymes via the p-ERK/p-CREB pathway in the fetal testis.

**Figure 7.**
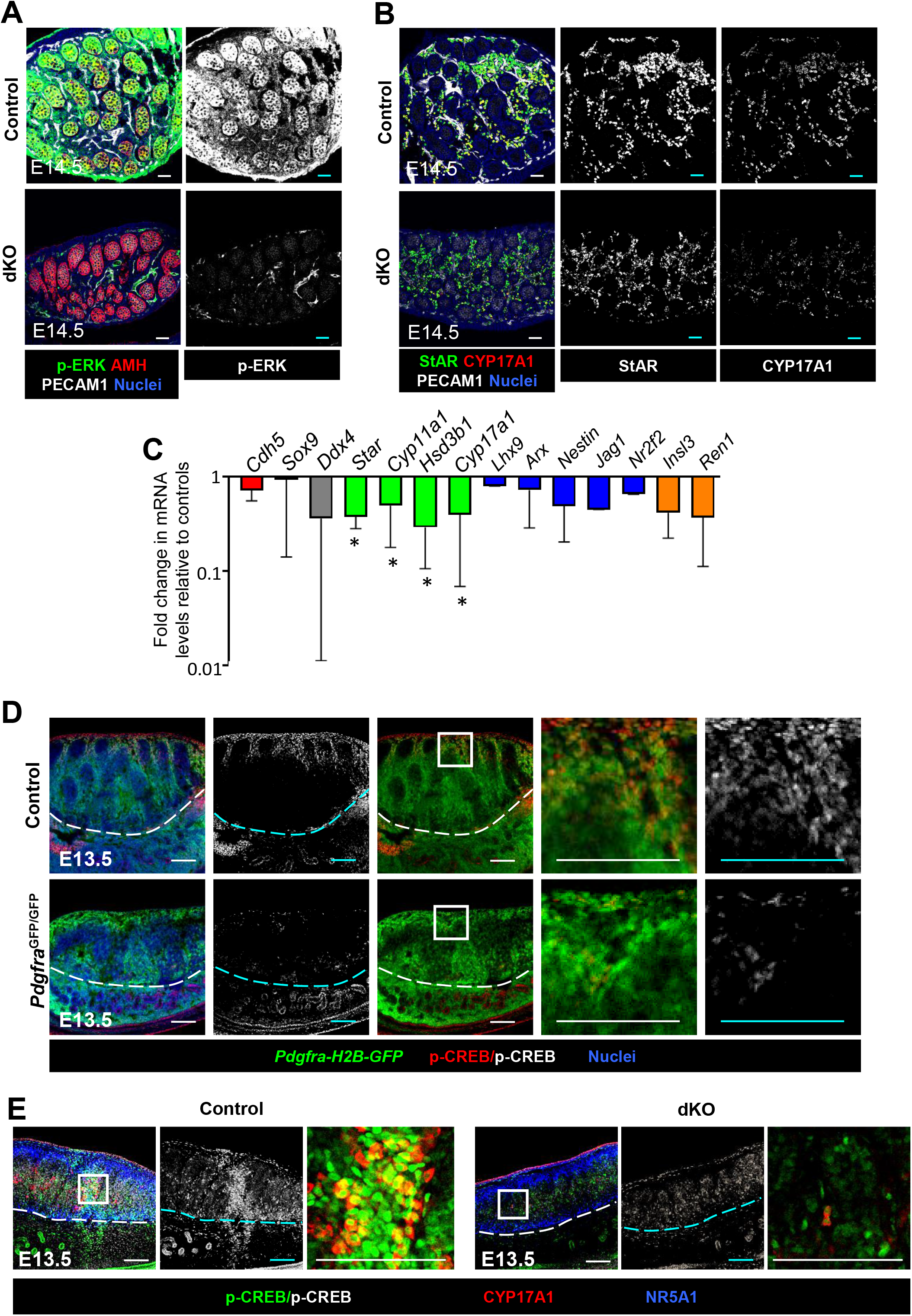
Steroidogenesis enzymes are disrupted after gonadal depletion of ERK signaling *in vivo*. (A-C) Immunofluorescence images (A,B) and qRT-PCR analyses (C) of E14.5 control (*Nr5a1*-Cre; *Mapk1^flox/+^*; *Mapk3^+/-^*) and dKO (*Nr5a1*-Cre; *Mapk1^flox/flox^*; *Mapk3^-/-^*) XY gonads. (D) Immunofluorescence images of E13.5 control and *Pdgfra*^GFP/GFP^ XY gonads. (E) Immunofluorescence images of E13.5 control (*Nr5a1*-Cre; *Mapk1^flox/+^*; *Mapk3^+/-^*) and dKO (*Nr5a1*-Cre; *Mapk1^flox/flox^*; *Mapk3^-/-^*) XY gonads. Scale bar: 100 μm.

## Discussion

*Pdgfra* has been shown to regulate testis cord morphogenesis and development of FLCs in XY gonads^9^. However, the mechanisms underlying how *Pdgfra* drives these processes are unclear. Here we investigated how cell migration might be involved in mediating the *Pdgfra* null testis phenotype. Mesonephric cell migration takes place between E11.5–16.5 in the XY mouse gonad^33^, and mesonephros-derived endothelial cells drive the formation of the coelomic artery and organization of testis cords. Previous work showed that knockout of *Pdgfra* in XY gonads impaired the migration of mesonephric cells, resulting in irregular testis cords^9^. Here we observed a reduction of *Cdh5* mRNA expression and a delay of vascularization in *Pdgfra*^GFP/GFP^ XY KI gonads, suggesting that a delay in vascularization caused by defective mesonephric cell migration might be responsible for irregular testis cords in KI gonads. Cell migration could be mediated via PDGFs through several downstream signaling pathways, including phosphoinositide-3-kinases (PI3K), MAPK, and ERK, which influence cell mobility, growth, and differentiation^34–37^. In particular, ERK plays a conserved role in driving cell migration in various systems^38–40^. The MAPK-specific inhibitors PD98059 and PI3K inhibitor LY294002, which reduce testis cord number or disrupt testis cord formation, respectively, reduce the protein level of p-ERK in the testis *ex vivo*, but the expression pattern and roles of ERK in the early fetal XY gonad have not been rigorously assessed. Here, we observed that p-ERK was broadly expressed in endothelial, interstitial, and Sertoli cells of 13.5 XY control gonads, while it was significantly reduced in *Pdgfra*^GFP/GFP^ XY KI gonads, indicating that PDGFRA activates the ERK pathway in the XY gonad. Our *ex vivo* results show that the ERK inhibitor U0126 blocked cord formation in E11.5 XY gonads but not in E12.5 XY gonads, consistent with our previous finding that vasculature is critical for initial testis cord formation at E11.5, but not for later maintenance of cord structure once already established^26^. We found that *Cdh5* mRNA expression was significantly reduced in both E11.5 and E12.5 U0126-treated XY gonads. These data suggest the possibility that U0126 hindered cord formation via disrupting vasculature in E11.5 XY gonads.

Early Growth Response 1 (EGR1), an early response transcription factor, is important for the initiation of growth and migration of vascular endothelial cells and mammary cells^10, 11^. EGR1 is activated by growth factors, cytokines, stress signals, and MAPK pathways including the ERK pathway^41, 42^. EGR1 promotes angiogenesis in a variety of contexts, including wound healing, corneal neovascularization, and tumor angiogenesis^43^. Neovascularization is impaired in subcutaneous Matrigel implants of *Egr1*-null mice^44^. The reduction of EGR1 in *Pdgfra*^GFP/GFP^ XY KI gonads suggest that EGR1 might be involved in vascular development of XY gonads. *EGR1*-knockout human vascular endothelial cells exhibit slower rates of proliferation and migration^45^, and here our analyses revealed that *Egr1*-knockdown mouse gonadal and mesonephric cells showed significantly compromised cell migration. Taken together, disruptions in vascular development in *Pdgfra*^GFP/GFP^ XY KI gonads might be due to a deficiency in *Egr1*-mediated cell migration. *Egr1* is a well-known ERK target gene in NIH/3T3 cells, which is a fibroblast cell line^46^. Our findings indicate that U0126 treatment impaired vascular development in XY gonads and disrupted cell migration in gonadal and mesonephric cells *ex vivo*, suggesting that active ERK signaling promotes vascular development via induction of cell migration. Additionally, both mRNA and protein levels of *Egr1* were significantly decreased in U0126-treated XY gonads and gonadal and mesonephric cells compared to controls, demonstrating that expression of *Egr1* was regulated by ERK activity in XY gonads. It has been reported that the ERK/EGR1 pathway is required for cell migration in mammary epithelial cells^10^. Our results suggest that ERK controls testis cord development via EGR1-mediated cell migration.

Vasculature is critical for the maintenance of FLC progenitors in the fetal testis^26^. Its disruption induces the differentiation of perivascular Nestin+ FLC progenitors into FLCs through inactivating vasculature-dependent Notch activity. However, defects in vascular development did not induce excessive differentiation of FLC progenitors in *Pdgfra*-null XY gonads, but instead was associated with a reduction in the number of FLCs. A previous study showed that coelomic epithelial proliferation was reduced in *Pdgfra*-knockout XY gonads, suggesting that the reduction of FLCs is due to defects in the proliferation of mesenchymal cells^9^. We found that the absence of *Pdgfra* impaired the proliferation of FLC progenitors at E12.5, but not at E13.5. Since the number of FLC progenitors was normal in E13.5 *Pdgfra* null XY gonads, we examined whether FLC progenitors failed to differentiation into FLCs due to the absence of *Pdgfra*. Interestingly, the number of NR5A1-bright cells, indicative of initial LC differentiation, in *Pdgfra*-knockout XY gonads was comparable to controls. We also observed some NR2F2-positive cells undergoing the transition to NR5A1+ cells, demonstrating that these progenitors were able to give rise to FLCs. However, most NR5A1 bright cells did not strongly express the steroidogenesis marker CYP11A1 in *Pdgfra-*null XY gonads, and a significantly increased number of cells did not express steroidogenesis markers at all. Taken together, our data shows that *Pdgfra* promotes the proliferation of mesenchymal FLC progenitors between E11.5-E12.5 and regulates the expression of steroidogenic genes in FLCs but is not essential for initial commitment to the FLC fate.

The MEK/ERK cascade in LCs is critical for maintenance of a functional population of ALCs in the adult testis, as deletion of *Mek1* and *Mek2* (official names *Map2k1* and *Map2k2*) in *Cyp17a1*-positive LCs resulted in a decreased number of CYP11A1-positive ALCs^18^. Furthermore, in the adult testis, PDGF-AA is able to activate ERK signaling in ALCs *in vitro*^47^, suggesting that PDGFA is the critical ligand. In *Pdgfra*-null XY gonads, the activation of ERK was reduced in endothelial, interstitial, and Sertoli cells. Therefore, we conditionally deleted *Erk2* in *Nr5a1*-expressing gonadal cells in the context of a global *Erk1* deletion using *Nr5a1*-Cre; *Mapk1^flox/flox^*; *Mapk3*^-/-^ embryos to explore whether *Pdgfra* regulates the expression of steroidogenic genes in FLCs via the ERK pathway. Knockout of the *Erk* genes significantly reduced the expression of steroidogenic genes but had no effect on the number of FLC progenitors and early-stage NR5A1+ FLCs, suggesting that *Pdgfra* controls the expression of downstream steroidogenic genes via the ERK pathway in XY gonads. The results of our promoter analyses using the JASPAR CORE collection of transcription factor binding sites showed that CREB has binding sites in the promoter regions of *Star*, *Cyp11a1*, *Hsd3b1*, and *Cyp17a1*. We also observed a reduction of p-CREB staining in the interstitial cells of *Pdgfra*- and *Erk*-knockout XY gonads. As a result, we propose that *Pdgfra* is critical for the expression of steroidogenic genes in FLCs via an ERK/p-CREB axis in the fetal testis.

Overall, our studies have provided insights into the distinct roles of *Pdgfra* in testis cord formation and FLC development during fetal stages. *Pdgfra* drives cell migration and organizes testis cords via the ERK/EGR1 pathway and regulates the expression of steroidogenic genes in FLCs via an ERK/p-CREB mechanism. The initial mechanisms and integration of these pathways provide a starting point for future research and will help us understand the underlying mechanisms that drive fetal testis development.

## Methods

### Animals

*Pdgfra*-H2B-eGFP (KI) mice (JAX stock #007669), *Nr5a1*-Cre mice (JAX stock #012462), and Cre-responsive *Rosa*-Tomato mice (JAX stock #007914), were obtained from The Jackson Laboratory. Given the embryonic lethality previously reported for *Pdgfra^GFP/GFP^* KI embryos^29^, which is potentially attributed to genetic background, we generated F1 *Pdgfra* heterozygous mice by crossing *Pdgfra* heterozygous males (maintained on the original C57BL/6J background) with CD-1 (Charles River) females to improve the recovery rate of homozygous mutant KI gonads (Fig. S1). All timed matings were between two F1 *Pdgfra^GFP/+^*heterozygous parents. *Kdr-*myr-mCherry mice^48^ were obtained from Dr. Mary Dickinson. *Mapk1^flox/flox^*; *Mapk3^-/-^* mice^49, 50^ were obtained from Dr. Jeffery Molkentin. Timed matings were checked daily for the presence of a vaginal plug. When plugs were detected in the morning, embryos were considered to be E0.5 at 12:00 noon. All mice were housed in accordance with National Institutes of Health guidelines, and experimental protocols were approved by the Institutional Animal Care and Use Committee (IACUC) of Cincinnati Children’s Hospital Medical Center (animal experimental protocol number IACUC2021-0016).

### Immunofluorescence

Tissues were fixed in 4% paraformaldehyde (PFA) with 0.1% Triton X-100 overnight at 4°C. Whole-mount immunofluorescence was performed on gonads at stages E11.5, E12.5, and E13.5. After several washes in PBTx (PBS + 0.1% Triton X-100), samples were incubated in blocking solution containing 10% FBS and 3% bovine serum albumin (BSA) for 1 h at room temperature. Gonads were incubated with primary antibodies (diluted in blocking solution) on a rocker overnight at 4°C. After several washes in PBTx, fluorescent secondary antibodies were applied for 4 h rocking at room temperature. After several washes in PBTx, samples were mounted on slides in Fluoromount-G (SouthernBiotech). Primary antibodies and dilutions used in this study are listed in Supplementary Table S2. *Kdr-*myr-mCherry and *Pdgfra*-H2B-eGFP expression were detected via endogenous fluorescence. Alexa-488-, Alexa-555-, and Alexa-647-conjugated secondary antibodies (Molecular Probes/Life Technologies/Thermo Fisher) were all used at 1:500. Nuclear staining was performed using 2 μg/mL Hoechst 33342 (#H1399, Thermo Fisher), and Hoechst staining is labeled as “Nuclei” in all figures. Pictures were taken on a Nikon A1 inverted LUNV microscope (Nikon, Tokyo, Japan) equipped with NIS-Elements imaging software (Nikon). At least two independent experiments were performed and within each experiment multiple XY gonads (*n* = 3-5) were used.

### Live Imaging

All imaging experiments were performed on a Zeiss LSM510 or LSM710 as previously described^51^. Z-stacks were collected every 10-15 minutes. All movies are maximum intensity projections at each time point, unless otherwise noted.

### Gonad Culture

E11.5 and E12.5 XY gonads were cultured *ex vivo* using an agar culture method as previously described^52^. Gonads were cultured on 1.5% agarose gel in Dulbecco’s Minimal Eagle Medium (DMEM) supplemented with 10% FBS and 1% penicillin–streptomycin at 37°C and 5% CO_2_. Medium change was performed every 24 hours. U0126, a selective inhibitor of MAPK/ERK kinases, was dissolved in DMSO and used at a final concentration of 50 μM, accounting for the volume of the agar block. Gonads were harvested for immunofluorescence or RNA extraction after 48 hours. At least three independent experiments were performed and within each experiment multiple XY gonads (*n* = 3-5) were used.

### RNA extraction and qRT-PCR (quantitative reverse transcriptase PCR)

Gonads were separated from the mesonephros, and total RNA was extracted using a standard TRIzol reagent (Invitrogen/Thermo Fisher) protocol. Reverse transcription reactions were performed with 500 ng of total RNA, using an iScript cDNA synthesis kit (Bio-Rad). qRT-PCR was performed on the StepOnePlus Real-Time PCR System (Applied Biosystems/Thermo Fisher) using the Fast SYBR Green Master Mix (Applied Biosystems/Thermo Fisher). Cycling conditions were: 95°C for 20 s, followed by 40 cycles of 95°C for 3 s and 60°C for 30 s. Primer specificity for a single amplicon was verified by melt curve analysis or agarose gel electrophoresis. All reactions were run in triplicate. *Gapdh* was used for normalization. Data from qRT-PCR was calculated relative to controls using the ΔΔCt method. Results were shown as mean ± SD. At least 3 biological replicates were used per condition for all experiments, and each biological replicate contained at least 2 male gonads. A two-tailed Student t-test was performed to calculate *P* values, in which *P*<0.05 was considered statistically significant. Statistical analyses were performed using Prism version 5.0 (GraphPad). All primer sequences used for qRT-PCR are listed in Supplementary Table S3.

### Cell isolation and siRNA transfection

Gonad and mesonephroi from E12.5 fetal testes were dissociated with 0.05% trypsin (Sigma) for 5 minutes at 37°C. DMEM medium containing 10% FBS was used to inhibit trypsin activity. The supernatants were filtrated through a 70μm cell strainer and centrifuged to collect the gonadal and mesonephric cells. The isolated cells were grown in DMEM medium containing 10% FBS and 1% penicillin–streptomycin at 37°C and 5% CO_2_. The medium was changed to remove the unattached cells after an initial 1-hour culture and allowed to attach overnight.

Gonadal and mesonephric cells were transfected with siRNA (30nM) by Lipofectamine 2000 (Invitrogen, Carlsbad, CA) according to the manufacturer’s instructions. Negative control (Assay ID: 4390843) and *Egr1* siRNAs (Assay ID: 157282) were purchased from Thermo Fisher Scientific. Cells were collected 2 days after treatment and RNA was extracted for qRT-PCR analyses.

### Transwell assay

Migration of gonadal and mesonephric cells was analyzed using a 6.5mm Transwell with 8.0 μm Pore Polyester Membrane Insert (3464, Corning, Somerville, MA). Equal numbers of gonadal and mesonephric cells were seeded in individual chambers. After siRNA transfection, DMEM medium with 2% or 10% FBS was used for upper and lower chambers, respectively. 2 days after siRNA transfection, cells remaining on the top of the membrane were removed by cotton-tipped applicators. The migrated cells were stained with crystal violet and photographed using an EVOS cell imaging system (Thermo Fisher Scientific). Migrated cells were quantified using ImageJ software (NIH).

### Statistics

All cell count charts for cell proliferation represent manual counts of pHH3+ cells in the coelomic domain. The coelomic domain was defined as the region extending from the top of testis cords to the coelomic surface. For testis cord counts, the number of testis cords was manually counted from 10 control XY gonads and 9 *Pdgfra*^GFP/GFP^ XY KI gonads. Error bars indicate +/- SEM and significance was calculated using an independent two-tailed Student t-test with *P*<0.05 considered statistically significant.

## References

1. Chen, D., Zuo, D., Luan, C., Liu, M., Na, M., Ran, L., Sun, Y., Persson, A., Englund, E., Salford, L.G., et al. (2014). Glioma cell proliferation controlled by ERK activity dependent surface expression of PDGFRA. PLoS One 9, e87281. 10.1371/journal.pone.0087281.

2. Hoch, R.V., and Soriano, P. (2003). Roles of PDGF in animal development. Development 130, 4769–4784. 10.1242/dev.00721.

3. Leveen, P., Pekny, M., Gebre-Medhin, S., Swolin, B., Larsson, E., and Betsholtz, C. (1994). Mice deficient for PDGF B show renal, cardiovascular, and hematological abnormalities. Genes Dev 8, 1875–1887. 10.1101/gad.8.16.1875.

4. Soriano, P. (1994). Abnormal kidney development and hematological disorders in PDGF beta-receptor mutant mice. Genes Dev 8, 1888–1896. 10.1101/gad.8.16.1888.

5. Karlsson, L., Lindahl, P., Heath, J.K., and Betsholtz, C. (2000). Abnormal gastrointestinal development in PDGF-A and PDGFR-(alpha) deficient mice implicates a novel mesenchymal structure with putative instructive properties in villus morphogenesis. Development 127, 3457–3466. 10.1242/dev.127.16.3457.

6. Karlsson, L., Bondjers, C., and Betsholtz, C. (1999). Roles for PDGF-A and sonic hedgehog in development of mesenchymal components of the hair follicle. Development 126, 2611–2621. 10.1242/dev.126.12.2611.

7. Fruttiger, M., Karlsson, L., Hall, A.C., Abramsson, A., Calver, A.R., Bostrom, H., Willetts, K., Bertold, C.H., Heath, J.K., Betsholtz, C., and Richardson, W.D. (1999). Defective oligodendrocyte development and severe hypomyelination in PDGF-A knockout mice. Development 126, 457–467. 10.1242/dev.126.3.457.

8. Gnessi, L., Basciani, S., Mariani, S., Arizzi, M., Spera, G., Wang, C., Bondjers, C., Karlsson, L., and Betsholtz, C. (2000). Leydig cell loss and spermatogenic arrest in platelet-derived growth factor (PDGF)-A-deficient mice. J Cell Biol 149, 1019–1026. 10.1083/jcb.149.5.1019.

9. Brennan, J., Tilmann, C., and Capel, B. (2003). Pdgfr-alpha mediates testis cord organization and fetal Leydig cell development in the XY gonad. Genes Dev 17, 800–810. 10.1101/gad.1052503.

10. Tarcic, G., Avraham, R., Pines, G., Amit, I., Shay, T., Lu, Y., Zwang, Y., Katz, M., Ben-Chetrit, N., Jacob-Hirsch, J., et al. (2012). EGR1 and the ERK-ERF axis drive mammary cell migration in response to EGF. FASEB J 26, 1582–1592. 10.1096/fj.11-194654.

11. Wang, B., Guo, H., Yu, H., Chen, Y., Xu, H., and Zhao, G. (2021). The Role of the Transcription Factor EGR1 in Cancer. Front Oncol 11, 642547. 10.3389/fonc.2021.642547.

12. Sysol, J.R., Natarajan, V., and Machado, R.F. (2016). PDGF induces SphK1 expression via Egr-1 to promote pulmonary artery smooth muscle cell proliferation. Am J Physiol Cell Physiol 310, C983–992. 10.1152/ajpcell.00059.2016.

13. Li, M.W., Mruk, D.D., and Cheng, C.Y. (2009). Mitogen-activated protein kinases in male reproductive function. Trends Mol Med 15, 159–168. 10.1016/j.molmed.2009.02.002.

14. Mebratu, Y., and Tesfaigzi, Y. (2009). How ERK1/2 activation controls cell proliferation and cell death: Is subcellular localization the answer? Cell Cycle 8, 1168–1175. 10.4161/cc.8.8.8147.

15. Azami, T., Bassalert, C., Allegre, N., Valverde Estrella, L., Pouchin, P., Ema, M., and Chazaud, C. (2019). Regulation of the ERK signalling pathway in the developing mouse blastocyst. Development 146. 10.1242/dev.177139.

16. Guo, Y.J., Pan, W.W., Liu, S.B., Shen, Z.F., Xu, Y., and Hu, L.L. (2020). ERK/MAPK signalling pathway and tumorigenesis. Exp Ther Med 19, 1997–2007. 10.3892/etm.2020.8454.

17. Shupe, J., Cheng, J., Puri, P., Kostereva, N., and Walker, W.H. (2011). Regulation of Sertoli-germ cell adhesion and sperm release by FSH and nonclassical testosterone signaling. Mol Endocrinol 25, 238–252. 10.1210/me.2010-0030.

18. Yamashita, S., Tai, P., Charron, J., Ko, C., and Ascoli, M. (2011). The Leydig cell MEK/ERK pathway is critical for maintaining a functional population of adult Leydig cells and for fertility. Mol Endocrinol 25, 1211–1222. 10.1210/me.2011-0059.

19. Uzumcu, M., Westfall, S.D., Dirks, K.A., and Skinner, M.K. (2002). Embryonic testis cord formation and mesonephric cell migration requires the phosphotidylinositol 3 kinase signaling pathway. Biol Reprod 67, 1927–1935. 10.1095/biolreprod.102.006254.

20. Chen, S.R., and Liu, Y.X. (2016). Testis Cord Maintenance in Mouse Embryos: Genes and Signaling. Biol Reprod 94, 42. 10.1095/biolreprod.115.137117.

21. Nassar, G.N., and Leslie, S.W. (2022). Physiology, Testosterone. In StatPearls.

22. Griswold, S.L., and Behringer, R.R. (2009). Fetal Leydig cell origin and development. Sex Dev 3, 1–15. 10.1159/000200077.

23. Shima, Y., Miyabayashi, K., Haraguchi, S., Arakawa, T., Otake, H., Baba, T., Matsuzaki, S., Shishido, Y., Akiyama, H., Tachibana, T., et al. (2013). Contribution of Leydig and Sertoli cells to testosterone production in mouse fetal testes. Mol Endocrinol 27, 63–73. 10.1210/me.2012-1256.

24. O’Shaughnessy, P.J., Baker, P.J., Heikkila, M., Vainio, S., and McMahon, A.P. (2000). Localization of 17beta-hydroxysteroid dehydrogenase/17-ketosteroid reductase isoform expression in the developing mouse testis--androstenedione is the major androgen secreted by fetal/neonatal leydig cells. Endocrinology 141, 2631–2637. 10.1210/endo.141.7.7545.

25. DeFalco, T., Takahashi, S., and Capel, B. (2011). Two distinct origins for Leydig cell progenitors in the fetal testis. Dev Biol 352, 14–26. 10.1016/j.ydbio.2011.01.011.

26. Kumar, D.L., and DeFalco, T. (2018). A perivascular niche for multipotent progenitors in the fetal testis. Nat Commun 9, 4519. 10.1038/s41467-018-06996-3.

27. Stevant, I., Neirijnck, Y., Borel, C., Escoffier, J., Smith, L.B., Antonarakis, S.E., Dermitzakis, E.T., and Nef, S. (2018). Deciphering Cell Lineage Specification during Male Sex Determination with Single-Cell RNA Sequencing. Cell Rep 22, 1589–1599. 10.1016/j.celrep.2018.01.043.

28. Schmahl, J., Rizzolo, K., and Soriano, P. (2008). The PDGF signaling pathway controls multiple steroid-producing lineages. Genes Dev 22, 3255–3267. 10.1101/gad.1723908.

29. Hamilton, T.G., Klinghoffer, R.A., Corrin, P.D., and Soriano, P. (2003). Evolutionary divergence of platelet-derived growth factor alpha receptor signaling mechanisms. Mol Cell Biol 23, 4013–4025. 10.1128/MCB.23.11.4013-4025.2003.

30. Combes, A.N., Wilhelm, D., Davidson, T., Dejana, E., Harley, V., Sinclair, A., and Koopman, P. (2009). Endothelial cell migration directs testis cord formation. Dev Biol 326, 112–120. 10.1016/j.ydbio.2008.10.040.

31. Kaetzel, D.M. (2003). Transcription of the platelet-derived growth factor A-chain gene. Cytokine Growth Factor Rev 14, 427–446. 10.1016/s1359-6101(03)00051-0.

32. Namura, S., Iihara, K., Takami, S., Nagata, I., Kikuchi, H., Matsushita, K., Moskowitz, M.A., Bonventre, J.V., and Alessandrini, A. (2001). Intravenous administration of MEK inhibitor U0126 affords brain protection against forebrain ischemia and focal cerebral ischemia. Proc Natl Acad Sci U S A 98, 11569–11574. 10.1073/pnas.181213498.

33. Martineau, J., Nordqvist, K., Tilmann, C., Lovell-Badge, R., and Capel, B. (1997). Male-specific cell migration into the developing gonad. Curr Biol 7, 958–968. 10.1016/s0960-9822(06)00415-5.

34. Tothova, Z., Semelakova, M., Solarova, Z., Tomc, J., Debeljak, N., and Solar, P. (2021). The Role of PI3K/AKT and MAPK Signaling Pathways in Erythropoietin Signalization. Int J Mol Sci 22. 10.3390/ijms22147682.

35. Dinsmore, C.J., and Soriano, P. (2018). MAPK and PI3K signaling: At the crossroads of neural crest development. Dev Biol 444 *Suppl 1*, S79–S97. 10.1016/j.ydbio.2018.02.003.

36. Singh, J., Sharma, K., Frost, E.E., and Pillai, P.P. (2019). Role of PDGF-A-Activated ERK Signaling Mediated FAK-Paxillin Interaction in Oligodendrocyte Progenitor Cell Migration. J Mol Neurosci 67, 564–573. 10.1007/s12031-019-1260-1.

37. Ying, H.Z., Chen, Q., Zhang, W.Y., Zhang, H.H., Ma, Y., Zhang, S.Z., Fang, J., and Yu, C.H. (2017). PDGF signaling pathway in hepatic fibrosis pathogenesis and therapeutics (Review). Mol Med Rep 16, 7879–7889. 10.3892/mmr.2017.7641.

38. Samson, S.C., Khan, A.M., and Mendoza, M.C. (2022). ERK signaling for cell migration and invasion. Front Mol Biosci 9, 998475. 10.3389/fmolb.2022.998475.

39. Samson, S.C., Elliott, A., Mueller, B.D., Kim, Y., Carney, K.R., Bergman, J.P., Blenis, J., and Mendoza, M.C. (2019). p90 ribosomal S6 kinase (RSK) phosphorylates myosin phosphatase and thereby controls edge dynamics during cell migration. J Biol Chem 294, 10846–10862. 10.1074/jbc.RA119.007431.

40. Li, J., Zhang, S., Soto, X., Woolner, S., and Amaya, E. (2013). ERK and phosphoinositide 3-kinase temporally coordinate different modes of actin-based motility during embryonic wound healing. J Cell Sci 126, 5005–5017. 10.1242/jcs.133421.

41. Shan, J., Dudenhausen, E., and Kilberg, M.S. (2019). Induction of early growth response gene 1 (EGR1) by endoplasmic reticulum stress is mediated by the extracellular regulated kinase (ERK) arm of the MAPK pathways. Biochim Biophys Acta Mol Cell Res 1866, 371–381. 10.1016/j.bbamcr.2018.09.009.

42. Khachigian, L.M. (2021). Early Growth Response-1, an Integrative Sensor in Cardiovascular and Inflammatory Disease. J Am Heart Assoc 10, e023539. 10.1161/JAHA.121.023539.

43. Fahmy, R.G., Dass, C.R., Sun, L.Q., Chesterman, C.N., and Khachigian, L.M. (2003). Transcription factor Egr-1 supports FGF-dependent angiogenesis during neovascularization and tumor growth. Nat Med 9, 1026–1032. 10.1038/nm905.

44. Khachigian, L.M. (2006). Early growth response-1 in cardiovascular pathobiology. Circ Res 98, 186–191. 10.1161/01.RES.0000200177.53882.c3.

45. Santiago, F.S., Li, Y., and Khachigian, L.M. (2021). Serine 26 in Early Growth Response-1 Is Critical for Endothelial Proliferation, Migration, and Network Formation. J Am Heart Assoc 10, e020521. 10.1161/JAHA.120.020521.

46. Torii, S., Kusakabe, M., Yamamoto, T., Maekawa, M., and Nishida, E. (2004). Sef is a spatial regulator for Ras/MAP kinase signaling. Dev Cell 7, 33–44. 10.1016/j.devcel.2004.05.019.

47. Tsai, Y.C., Kuo, T.N., Chao, Y.Y., Lee, P.R., Lin, R.C., Xiao, X.Y., Huang, B.M., and Wang, C.Y. (2022). PDGF-AA activates AKT and ERK signaling for testicular interstitial Leydig cell growth via primary cilia. J Cell Biochem. 10.1002/jcb.30345.

48. Poche, R.A., Larina, I.V., Scott, M.L., Saik, J.E., West, J.L., and Dickinson, M.E. (2009). The Flk1-myr::mCherry mouse as a useful reporter to characterize multiple aspects of ocular blood vessel development and disease. Dev Dyn 238, 2318–2326. 10.1002/dvdy.21886.

49. Samuels, I.S., Karlo, J.C., Faruzzi, A.N., Pickering, K., Herrup, K., Sweatt, J.D., Saitta, S.C., and Landreth, G.E. (2008). Deletion of ERK2 mitogen-activated protein kinase identifies its key roles in cortical neurogenesis and cognitive function. J Neurosci 28, 6983–6995. 10.1523/JNEUROSCI.0679-08.2008.

50. Pages, G., Guerin, S., Grall, D., Bonino, F., Smith, A., Anjuere, F., Auberger, P., and Pouyssegur, J. (1999). Defective thymocyte maturation in p44 MAP kinase (Erk 1) knockout mice. Science 286, 1374–1377. 10.1126/science.286.5443.1374.

51. Coveney, D., Cool, J., Oliver, T., and Capel, B. (2008). Four-dimensional analysis of vascularization during primary development of an organ, the gonad. Proc Natl Acad Sci U S A 105, 7212–7217. 0707674105 [pii] 10.1073/pnas.0707674105.

52. Li, S.Y., Bhandary, B., Gu, X., and DeFalco, T. (2022). Perivascular cells support folliculogenesis in the developing ovary. Proc Natl Acad Sci U S A 119, e2213026119. 10.1073/pnas.2213026119.

