## Supplementary Information for "Testis morphogenesis and fetal Leydig cell development are mediated through ERK signaling activated by platelet derived growth factor receptor alpha"

Contains:

Supplementary Figures S1-S5
Supplementary Figure Legends
Supplementary Tables S1-S3
Supplementary References

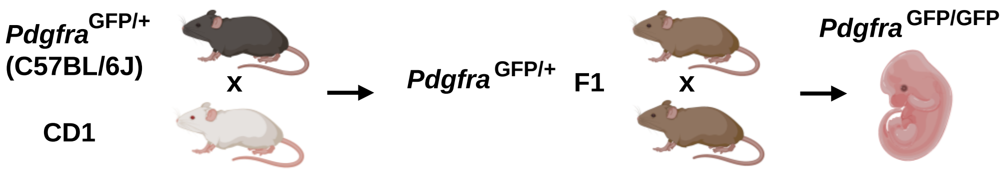


### Supplementary Figure S1

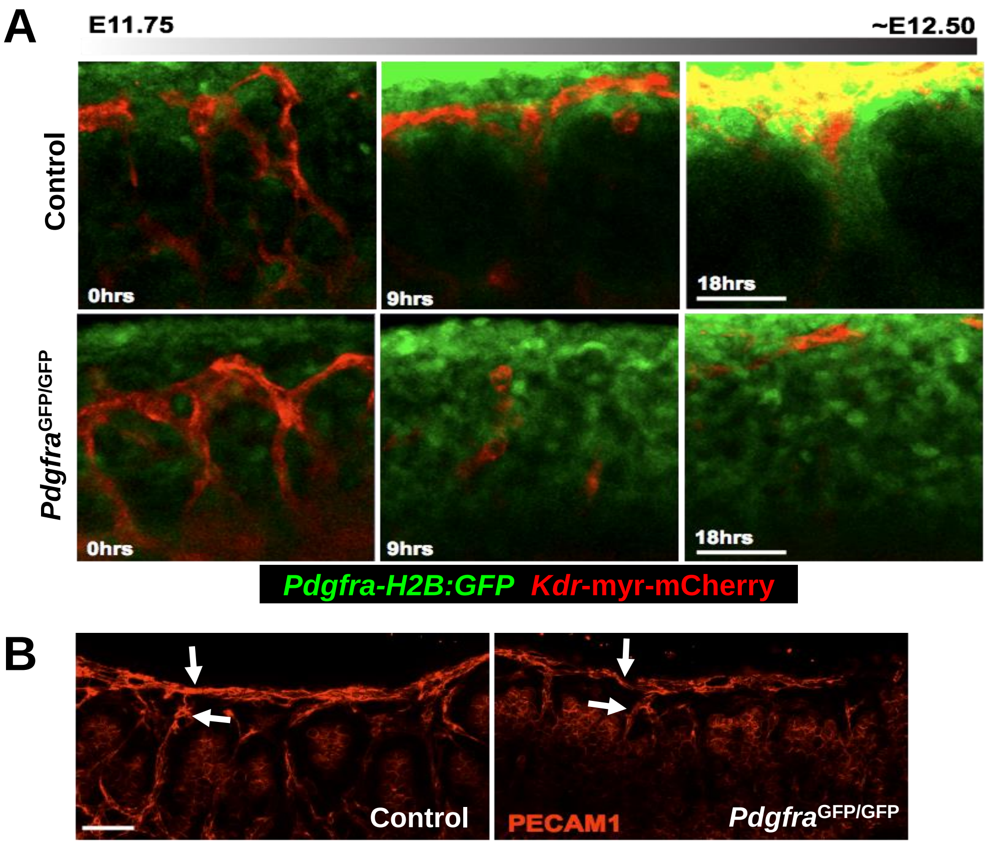


### Supplementary Figure S2

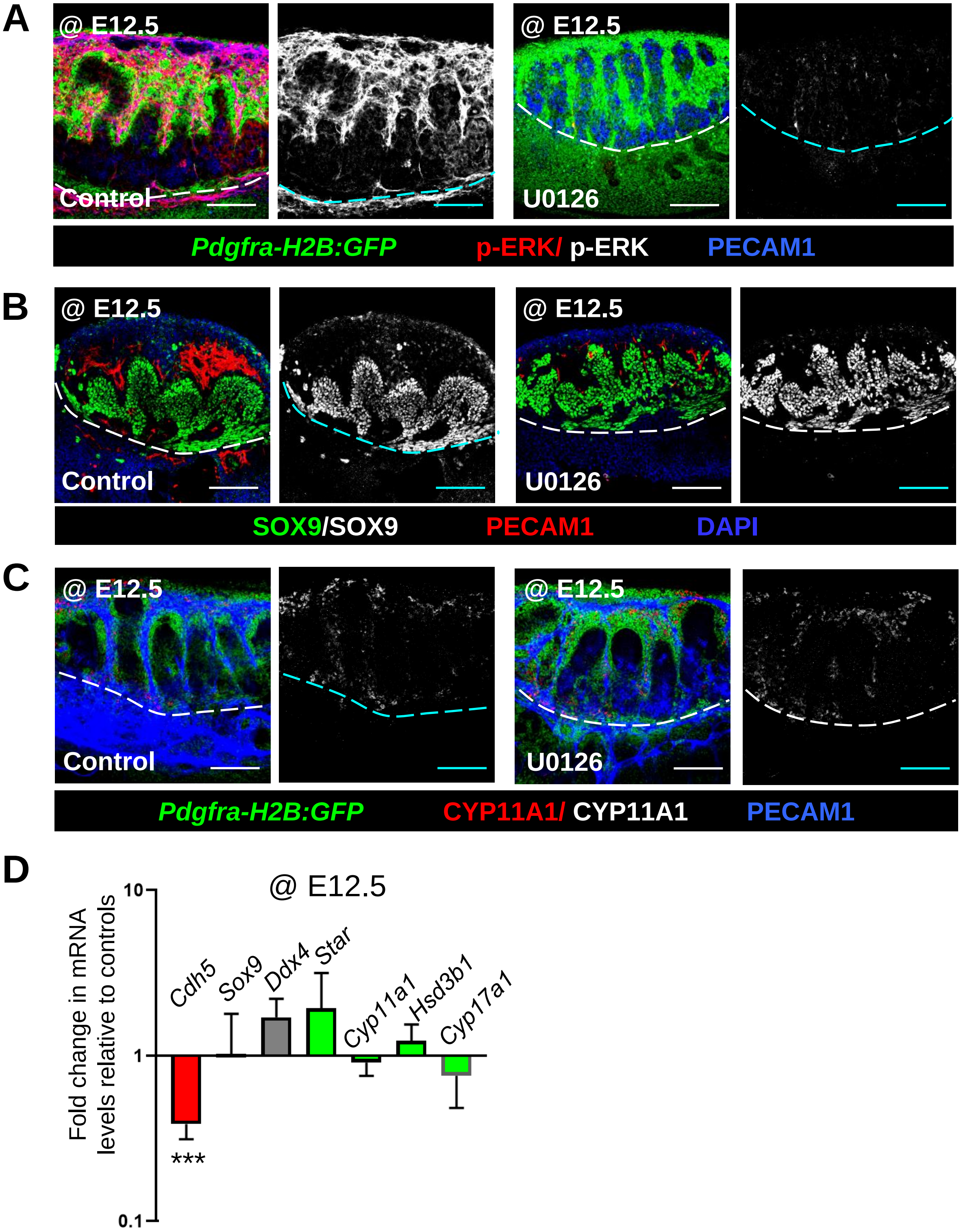


### Supplementary Figure S3

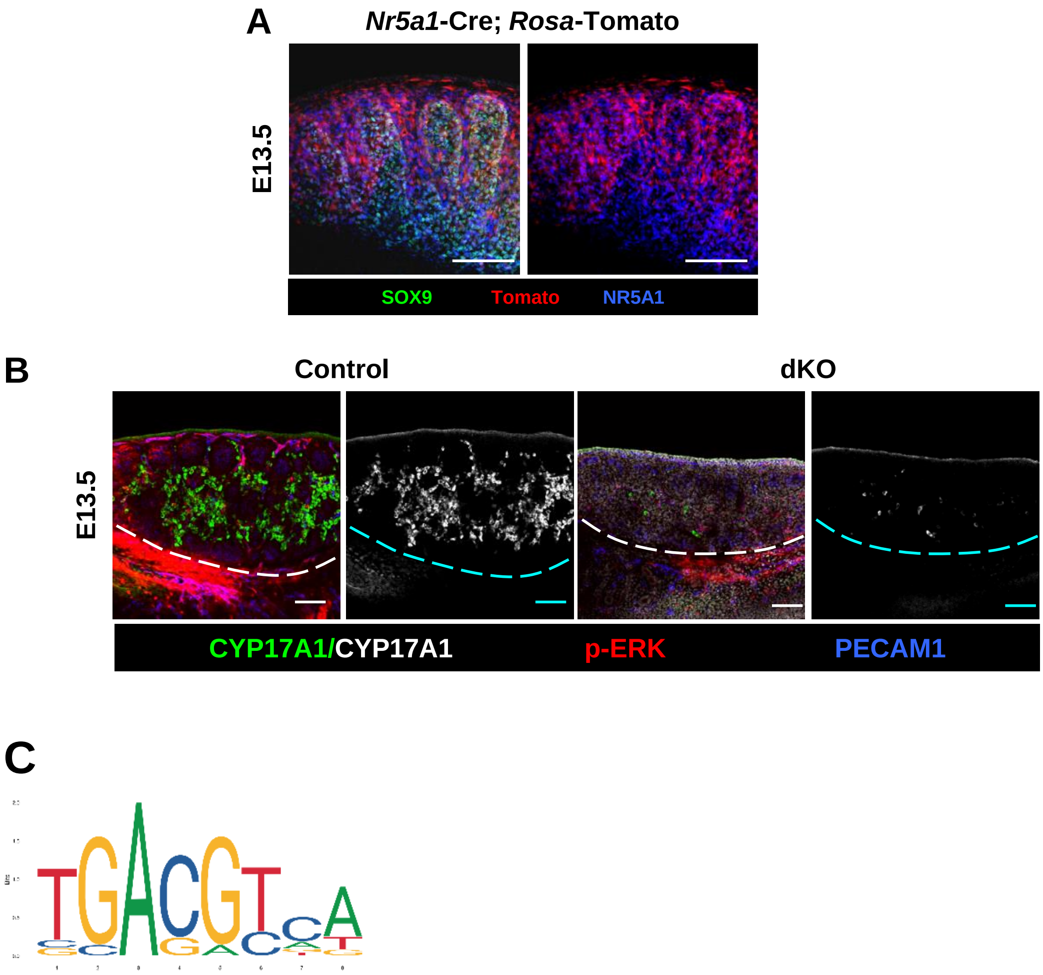


### Supplementary Figure S4

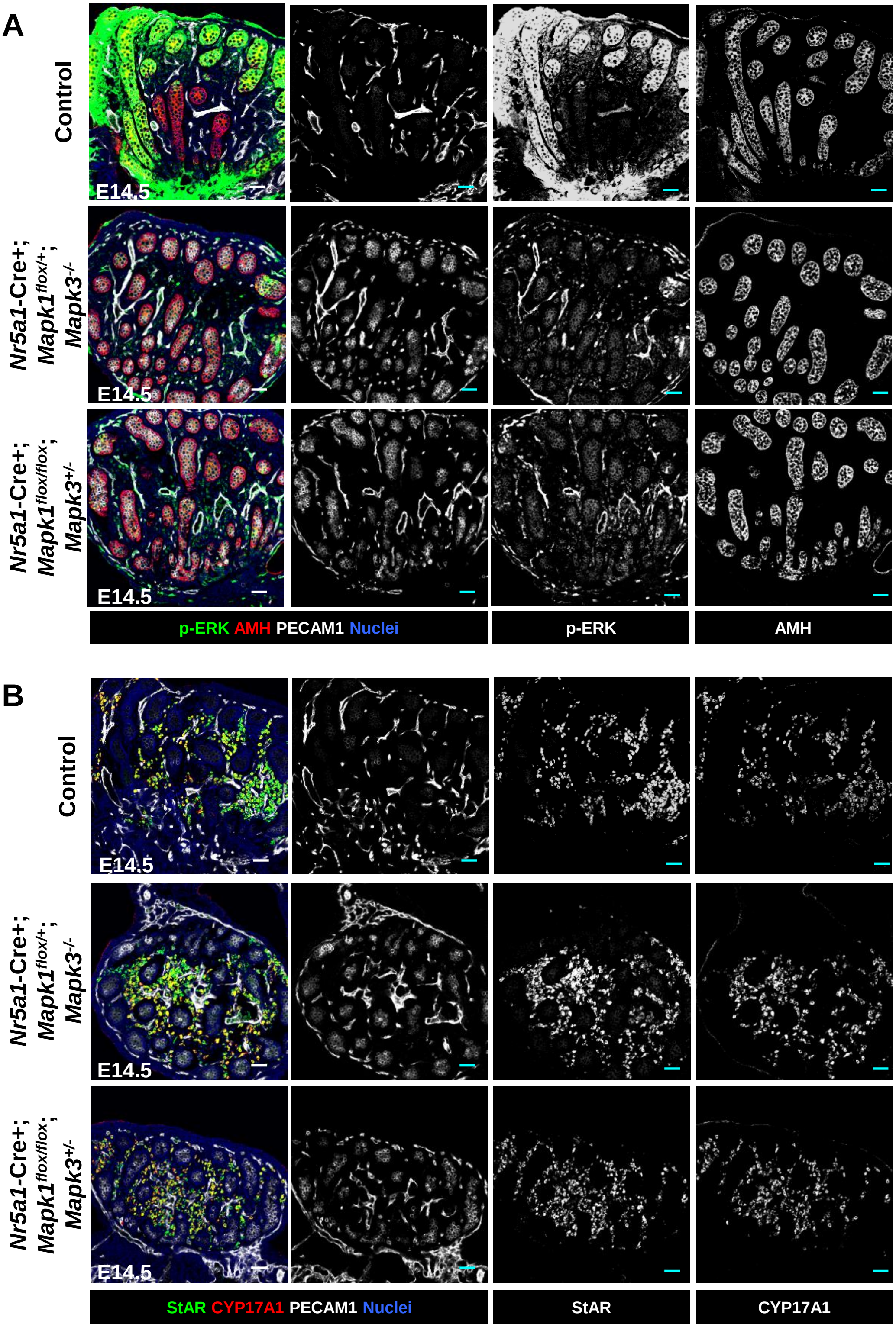


### Supplementary Figure S5

**Supplementary Figure Legends**

**Supplementary Figure S1. Breeding strategy to obtain *Pdgfra*$^{GFP/GFP}$ embryos.** Cartoon depicts experimental strategy to generate *Pdgfra*$^{GFP/GFP}$ KI embryos.

**Supplementary Figure S2. Endothelial cell migration is delayed in *Pdgfra*$^{GFP/GFP}$ KI XY gonads.**

(A) Live imaging of *Kdr*-myr-mCherry; *Pdgfra*$^{GFP/+}$ control (top) and *Kdr*-myr-mCherry; *Pdgfra*$^{GFP/GFP}$ (bottom) XY gonads initiated at E11.75. (C) Immunostaining of E12.5 XY gonads with PECAM1, which labels germ and endothelial cells. Arrowheads denote vascular branch points. Scale bar: 50 µm.

**Supplementary Figure S3. Inhibition of ERK signaling at E12.5 does not disrupt testis cord patterning.**

Immunofluorescence images (A-C) and qRT-PCR analyses (D) of E12.5 XY gonads cultured with U0126 for 48 hours *ex vivo*. Scale bar: 100 µm. ****P*<0.001.

**Supplementary Figure S4. Activity pattern of *Nr5a1*-Cre, steroidogenesis enzyme expression in ERK dKO XY gonads, and CREB binding motif.**

(A) E13.5 *Nr5a1*-Cre; *Rosa*-Tomato XY gonad, showing broad gonadal Tomato expression. (B) E13.5 control (*Nr5a1*-Cre+; *Mapk1*$^{flox/+}$; *Mapk3*$^{+/-}$) and dKO (*Nr5a1*-Cre+; *Mapk1*$^{flox/flox}$; *Mapk3*$^{-/-}$) XY gonads. Scale bar: 100 µm. (C) Binding motif of CREB transcription factor.

**Supplementary Figure S5. Steroidogenesis enzyme expression in E14.5 ERK sKO XY gonads.**

(A,B) Immunofluorescence images of E14.5 control (*Nr5a1*-Cre+; *Mapk1*$^{flox/+}$; *Mapk3*$^{+/-}$) and sKO (*Nr5a1*-Cre+; *Mapk1*$^{flox/+}$; *Mapk3*$^{-/-}$ or *Nr5a1*-Cre+; *Mapk1*$^{flox/flox}$; *Mapk3*$^{+/-}$) XY gonads. Scale bar: 100 µm.

**Table S1. Predicted binding sites of CREB.**

| Matrix ID | Transcription factor | Score | Relative score | Gene | Predicted sequence |
|---|---|---|---|---|---|
| MA0018.2 | CREB | 8.00 | 0.87 | *Star* | TGAAGTCA |
| | | 8.30 | 0.88 | *Cyp11a1* | TGAAGTCA |
| | | 9.93 | 0.94 | *Hsd3b1* | TGACGTCT |
| | | 8.16 | 0.88 | *Cyp17a1* | TGACGGCA |

**Table S2. List of primary antibodies used for immunofluorescence.**

| Primary Antibody | Dilution | Source/Reference |
|---|---|---|
| Goat anti-JAG1 | 1:1,000 | R&D #AF599 |
| Mouse anti-NR2F2 | 1:500 | Perseus Proteomics #PP-H7147-00 |
| Rabbit anti-CYP11A1 | 1:2,000 | D. Wilhelm[1] |
| Rabbit anti-Nestin | 1:1,000 | Biolegend #PRB-315C |
| Rabbit anti-SOX9 | 1:500 | EMD Millipore #AB5535 |
| Rat anti-PECAM1 | 1:250 | BD Pharmingen #553370 |
| Goat anti-PECAM1 | 1:250 | R&D Systems #AF3628 |
| Goat anti-CYP17A1 | 1:500 | Santa Cruz #sc-46081 |
| Mouse anti-AMH | 1:500 | Santa Cruz #sc-365643 |
| Rabbit anti-p-ERK | 1:2,000 | Cell Signal Technology #9101 |
| Rabbit anti-EGR1 | 1:2,000 | Cell Signal Technology #4153s |
| Rabbit anti-pHH3 | 1:500 | Millipore #AF4465-SP |
| Rat anti-NR5A1 | 1:300 | Cosmo Bio #KAL-KO610 |
| Rabbit anti-StAR | 1:500 | Cell Signal Technology #8449 |
| Rabbit anti-p-CREB | 1:2,000 | Cell Signal Technology #9198 |

**Table S3. Sequences of primers used for qRT-PCR analyses.**

| Gene name | Sequence (5' to 3') |
|---|---|
| *Arx* forward | CAAGGATGGTGAGGACAGC |
| *Arx* reverse | TCTGGAACCACACCTGGACT |
| *Cdh5* forward | TCCTCTGCATCCTCACTATCACA |
| *Cdh5* reverse | GTAAGTGACCAACTGCTCGTGAAT |
| *Cyp11a1* forward | TGGCCCCATTTACAGGGAGAA |
| *Cyp11a1* reverse | GGCATCTGAACTCTTAAACAGGA |
| *Cyp17a1* forward | CAGAGAAGTGCTCGTGAAGAAG |
| *Cyp17a1* reverse | AGGAGCTACTACTATCCGCAAA |
| *Ddx4* forward | TACTGTCAGACGCTCAACAGGA |
| *Ddx4* reverse | ATTCAACGTGTGCTTGCCCT |
| *Egr1* forward | GAGCGAACAACCCTATGAG |
| *Egr1* reverse | GTCGTTTGGCTGGGATAA |
| *Gapdh* forward | AGGTCGGTGTGAACGGATTTG |
| *Gapdh* reverse | TGTAGACCATGTAGTTGAGGTCA |
| *Hsd3b1* forward | CAAGTGTGCCAGCCTTCATCT |
| *Hsd3b1* reverse | TTCATGATTCTGTTCCTCGTGG |
| *Insl3* forward | CCTCCTGGCTATGTCATTGC |
| *Insl3* reverse | CCTGTGGTCCTTGCTTACTG |
| *Jag1* forward | TGACATGGATAAACACCAGCA |
| *Jag1* reverse | GCAGCCCACTGTCTGCTATAC |
| *Lhx9* forward | TGTAATGCCCCAAGATTTGTTCTCCC |
| *Lhx9* reverse | ACCAGCAGCCTTATCCACCTTCACAG |
| *Nestin* forward | GCTGGAACAGAGATTGGAAGG |
| *Nestin* reverse | CCAGGATCTGAGCGATCTGAC |
| *Nr2f2* forward | GCCATAGTCCTGTTCACCTC |
| *Nr2f2* reverse | GCTCCTAACGTACTCTTCCAAAG |
| *Pdgfrb* forward | TTCCAGGAGTGATACCAGCTT |
| *Pdgfrb* reverse | AGGGGGCGTGATGACTAGG |
| *Ren1* forward | CTCTCTGGGCACTCTTGTTGC |
| *Ren1* reverse | GGGAGGTAAGATTGGTCAAGGA |
| *Sox9* forward | GCGGAGCTCAGCAAGACTCTG |
| *Sox9* reverse | ATCGGGGTGGTCTTTCTTGTG |
| *Star* forward | TACATCCAGCAGGGAGAGGTG |
| *Star* reverse | CAGCGCACGCTCACGAAGTCT |

**Supplementary References**


1. Rastetter, R.H., Bernard, P., Palmer, J.S., Chassot, A.A., Chen, H., Western, P.S., Ramsay, R.G., Chaboissier, M.C., and Wilhelm, D. (2014). Marker genes identify three somatic cell types in the fetal mouse ovary. Dev Biol *394*, 242-252. 10.1016/j.ydbio.2014.08.013.
